# Hindbrain rhombomere centers harbor a heterogenous population of dividing progenitors which rely on Notch-signaling

**DOI:** 10.1101/2023.07.28.550945

**Authors:** Carla Belmonte-Mateos, Lydvina Meister, Cristina Pujades

## Abstract

Tissue growth and morphogenesis are interrelated processes, whose tight coordination is essential for the production of different cell fates and the timely precise allocation of stem cell capacities. The zebrafish embryonic brainstem, the hindbrain, exemplifies such coupling between spatiotemporal cell diversity acquisition and tissue growth, as the neurogenic commitment is differentially distributed over time. Here, we combined cell lineage and in vivo imaging approaches to reveal the emergence of different specific cell population properties within the very same rhombomeres, and focused on the hindbrain rhombomere centers. We studied the molecular identity of rhombomere centers, and showed that they harbor different progenitor capacities that change over time. By clonal analysis, we revealed that cells within the rhombomeres decrease the proliferative capacity over time to remain mainly in G1-phase. Proliferating progenitors give rise to neurons by asymmetric and symmetric neurogenic divisions, while maintaining the pool of progenitors. The proliferative capacity of these cells differs from their neighbors, and they are delayed in the onset of Notch-activity. By functional studies we demonstrated that they rely on Notch3-signaling to be maintained as non-committed progenitors. In this study we show that cells in rhombomere centers might share steps of a similar program, despite the neurogenic asynchrony from the rhombomere counterparts, to ensure proper tissue growth.

## INTRODUCTION

The generation of the brain cell diversity occurs concomitantly with tissue morphogenesis, resulting in changes in the position of neuronal progenitors and their derivatives over time. Thus, one of the main unsolved questions is how multiple cell types are generated and maintained in highly organized spatial patterns upon morphogenesis, and how changes in this ground plan can result in pathologies. The hindbrain has been proven a good experimental system to address the bases of neuronal diversity, since it displays stereotypic growth dynamics while undergoing tissue segmentation along the anteroposterior (AP) axis (Krumlauf and Wilkinson, 2021) and lumen formation (Lowery and Sive, 2004; Gutzman and Sive, 2010).

Hindbrain segmentation results in rhombomeres that constitute developmental units of gene expression and cell lineage compartments (Fraser et al., 1990; Lumsden, 2004; Jimenez-Guri et al., 2010; Krumlauf and Wilkinson, 2021), separated by interhombomeric boundaries. During the last years, it has been demonstrated that neurogenic capacities are sequentially allocated along the AP axis (Nikolaou et al., 2009; Esain et al., 2010; Gonzalez-Quevedo et al., 2010), with boundary cells engaging later than rhombomeres in neurogenesis (Peretz et al., 2016; Voltes et al., 2019; Pujades, 2020; Hevia et al., 2022). During the first neurogenic phase, hindbrain boundaries function as an elastic mesh to restrict cell intermingling between adjacent cell lineages (Calzolari et al., 2014; Letelier et al., 2018). Boundary cells undergo mechanical tension that activate YAP/TAZ-pathway that maintains them in the proliferative progenitor state (Voltes et al., 2019; Engel-Pizcueta and Pujades, 2021). Moreover, no synchronous patterning of neurogenesis is observed within the very same rhombomeres. Neurogenesis becomes confined to the cell population adjacent to rhombomere boundaries (Nikolaou et al., 2009) and the segment centers comprise a non-neurogenic progenitor population with a different molecular identity (Esain et al., 2010; Gonzalez-Quevedo et al., 2010; Tambalo et al., 2020), which signals through FGF to instruct neuronal organization (Gonzalez-Quevedo et al., 2010). Despite FGF-signaling is not essential for the survival or maintenance of hindbrain neural progenitors, it controls their fate by coordinately regulating *sox9* and *oligo2* (Esain et al., 2010). However, little is known about how these cells are maintained as progenitors.

Thus, in this study, we aimed at understanding how the neurogenic capacity was asynchronously distributed within the rhombomeres. We have explored their molecular identity and their proliferative capacity. Clonal analyses and cell proliferation experiments revealed that cells within the rhombomere centers are held as slow-dividing progenitors. Multicolor cell lineage analysis allowed us to demonstrate that rhombomere centers harbor cell that can engage in neurogenesis, contributing to the neuronal lineage. Functional analyses revealed the role of Notch3-signaling in the control of these cells as non-committed progenitors during hindbrain morphogenesis.

## MATERIALS AND METHODS

### Ethics declarations and approval for animal experiments

All procedures were approved by the institutional animal care and use ethic committee (Comitè Etica en Experimentació Animal, PRBB) and the Generalitat of Catalonia (Departament de Territori i Sostenibilitat), and implemented according to National and European regulations. Government and University veterinary-inspectors examine the animal facilities and procedures to endure that animal regulations are correctly followed. The PRBB animal house holds the AAALAC International approval B9900073. All the members entering the animal house have to hold the international FELASA accreditation. The Project License covering the proposed work (Ref 10642, GC) pays particular attention to the 3Rs.

### Zebrafish strains

Embryos were obtained by mating of adult fish using standard methods. All zebrafish strains were maintained individually as inbred lines. Tg[elA:GFP] (Labalette et al., 2011) and Mü4127 (Distel et al., 2009) transgenic lines were used as landmarks of rhombomeres 3 and 5, displaying GFP and DsRed respectively. Tg[BCP:H2AmCherry] and Tg[BCP:H2B-GFP] were used as landmark of hindbrain boundaries (Hevia et al., 2022). Tg[nestin:GFP] carries GFP at 3.9Kb upstream of the nestin promoter and labels neural progenitors (Lam et al., 2009). The Tg[HuC:GFP] line was used to label the whole neuronal differentiated domain (Park et al., 2000). The dual Fucci transgene, Tg[Fucci], ubiquitously produces both a Cerulean-tagged degron, which is detectable during the S/G2/M phases of the cell cycle, and a Cherry-tagged degron, which is only detectable during the G1 phase (Bouldin and Kimelman, 2014). The readout of Notch-activity line Tg[tp1:d2GFP] was built up using the *tp1* promoter (Clark et al., 2012; Hevia et al., 2022) and Notch-active cells express destabilized GFP. The *notch3^fh332^* null allele originated from ENU-induced nonsense mutation was identified by genotyping genomic DNA from fin clips or embryonic tails according to (Alunni et al., 2013). Embryos homozygous for *notch3^fh332/fh332^* were obtained by in-cross of heterozygous carriers. As controls, both *notch3^+/+^* and *notch3^fh332/+^* embryos were used since heterozygous embryos displayed wild type phenotype. The *hey1^ha11^* mutant allele harbors an 11bp deletion causing a frameshift leading to the production of a truncated protein; its presence was identified by genotyping genomic DNA from fin clips or embryonic tails according to (Than-Trong et al., 2018). Embryos homozygous for *hey1^ha11/ha11^* were obtained by in-cross of heterozygous carriers. As controls, both *hey^+/+^* and *hey1^ha11/+^* embryos were used since heterozygous embryos displayed wild type phenotype.

### Confocal imaging of whole-mount embryos

Anesthetized live embryos expressing genetically encoded fluorescence and stained fixed samples were mounted in 1% low melting point (LMP) agarose with the hindbrain positioned towards the glass-bottom of Petri dishes (Mattek) to achieve dorsal views of the hindbrain. Imaging was performed under a SP8 Leica confocal microscope.

### Whole-mount *in situ* hybridization

Embryo whole mount *in situ* hybridization was adapted from (Thisse and Thisse, 2008). The following riboprobes were generated by *in vitro* transcription from cloned cDNAs: *ascl1b* (Allende and Weinberg, 1994), *erm* (Riley et al., 2004), *meteorin* and *meteorin-like* (Tambalo et al., 2020), *neurod4* (Park et al., 2003), *neurog1* (Itoh and Chitnis, 2001), and *notch1b* (Dyer et al., 2014). The other probes were generated by PCR amplification adding the T7 or Sp6 promoter sequence in the Rv primers: *fabp7a* Fw: 5’ –GAC TGA ACT CAG CGA CTG TAC– 3’ and Rv: 5’ –AGG CCT CAA TAA TAC ACT CCC–T7 3’; *hey1* Fw: 5’ –GCA GAG ACT GCA CGT TAC CTC– 3’ and Rv: 5’ –GCC CCT ATT TCC ATG CTC CAG–T7 3’; *notch1a* Fw: 5’ –ACT TCG AAA TCG CTC CTC ATC– 3’ and Rv: 5’ –TCT TCC TGG AGA CGA CCA C–T7 3’; *notch3* Fw: 5’ –ATG GGG AAT TAC AGC CTT TG– 3’ and Rv: 5’ –GGC AAA CAA GCA ATT CGT A–SP6 3’; *slc1a2a* Fw: 5’ – GGG AAA GAT GGG AGA GAA GG– 3’ and Rv: 5’ –AGG ACT GTG TCT TGG CCA TC–T7 3’; *sox9b* Fw: 5’ –GGG CTG AAG ATG AGT GTG TC– 3’ and Rv: 5’ –CTT CAG ATC CGC TTA CTG CAC–T7 3’.

For fluorescent *in situ* hybridization, FLUO- and DIG-labeled probes were detected with TSA Fluorescein and Cy3, respectively.

### *In toto* embryo immunostainings

Embryos were blocked in 10% neutralized goat serum and 2% Bovine Serum Albumin (BSA) in PBST for 2h at RT, except after *in situ* hybridization where they were blocked in 5% neutralized goat serum in PBS-Tween20 (PBST) during 1h. They were then incubated O/N at 4°C with the corresponding primary antibody: anti-DsRed (1:500; Clontech), mouse anti-GFP ([1:100]; ThermoFisher), anti-fabp7a ([1:100], Millipore), rabbit anti-GFP ([1:500], TorreyPines), rabbit anti-pH3 (1:200; Upstate), rabbit anti-Sox2 ([1:100], Abcam), mouse anti-HuC ([1:100], ThermoFisher). After extensive washings with PBST, embryos were incubated with secondary antibodies conjugated with Alexa Fluor^®^488, Alexa Fluor^®^594, or Alexa Fluor^®^633 ([1:250], Invitrogen). DAPI ([1:5000], Molecular Probes) was used to label cell nuclei.

### Cell proliferation

#### EdU-incorporation experiments

Cells in S-phase were detected by EdU-incorporation using the Click-It^TM^ EdU Alexa Fluor^TM^ 647 Imaging Kit (C10340, Thermo Fisher Scientific) following the supplier instructions, with some modifications. Briefly, embryos were dechorionated, incubated in 500µM EdU diluted in 7% DMSO and placed on ice during the first hour for better EdU-incorporation. Afterwards, they were either washed three times with embryo medium to wash out the EdU or fixed in 4%PFA during 4 hours at RT and dehydrated in MetOH. After progressive rehydration, embryos were permeabilized with 10 mg/ml Proteinase K (Invitrogen) for 35min, post-fixed 40min in 4%PFA and washed in PBT. Embryos were then incubated for 1h in 1%DMSO/1% Triton X-100/PBS. The Click-iT reaction was carried out according to the manufacturer’s instructions for 60 min at RT, washed into PBT and then embryos were used for *in situ* hybridization with *fabp7a*. To determine the position of boundaries, and therefore rhombomeres centers, we used Tg[BCP:H2B-GFP] line and revealed the GFP by immunostaining. Imaging was undertaken with the AiryScan2 LSM980 using a 20× dry objective.

#### Cell cycle phase dynamics

In order to follow the cell cycle dynamics, the distribution of PCNA-GFP in the cell nuclei was assessed at different cell cycle phases. Mu4127 embryos, as landmark of r3 and r5, were injected between the 16-cell stage and 32-cell stage with PCNA-GFP mRNA, let to grow until the desired stage and *in vivo* imaged under the *AiryScan2* LSM980 microscope using 40× (1.2NA) glycerol immersive objective with the Zeiss acquisition software. Samples were maintained at 28°C while imaged. Optical sections were acquired through the entire hindbrain volume from dorsal to ventral (*z* = 1μm). For quantification, we counted the S-phase cells and the total number of PCNA-labelled nuclei in delimitated areas (76µm×7µm), both in the centers of rhombomeres 3, 4 and 5 and in the corresponding boundary flanking regions (n = 24 regions; N = 4 embryos). We then calculated the ratio between the number of nuclei in S-phase and the total number of PCNA-GFP labeled nuclei at 42 and at 48hpf. Finally, the percentage of S-phase cells both in boundary flanking regions and rhombomere centers was calculated for further comparisons.

### Zebrabow multicolor cell clonal analysis

For multicolor cell clonal analyses, Tg[HuC:GFP] embryos were injected with hsp:ZEBRABOW construct (Brockway et al., 2019), between 8-cell stage and 16-cell stage. Embryos were heat-shocked for 1h at 37°C just three hours before imaging. Then, they were washed in fresh embryo medium and kept at 28°C until the desired imaging time. Images were acquired with the Leica SP8 confocal microscope using a 20× water immersion objective with zoom ranging from 1.0 to 1.5. Embryos were then incubated at 28°C for 12 hours and reimaged with the same settings.

### TUNEL analyses

The distribution of apoptotic cells was determined by TdT-mediated dUTP nick-end labeling of the fragmented DNA (TUNEL, Roche). Briefly, embryos were fixed in 4% PFA, dehydrated in 100% MetOH, permeabilized with 80% acetone/H20, and preincubated with TUNEL mixture for 3h at 37°C according to the manufacturer’s instructions. DAPI ([1:5000], Molecular Probes) was used to label nuclei.

### Pharmacological treatments

Embryos were treated either with 10 μM of the gamma-secretase inhibitor LY411575 (Sigma-Aldrich) or DMSO for control. The treatment was applied to the fish water at 28.5°C for 6h at the indicated stages. After treatment, embryos were fixed in 4%PFA for further analysis.

### Sox2 and HuC volume quantifications

For the quantification of the progenitor and neuronal differentiation volumes in rhombomere 4 (Supplementary Figure 2), we developed a macro in ImageJ that can be found at https://github.com/cristinapujades/Belmonte-Mateos-et-al-2023

## RESULTS

### Cells within the rhombomere centers display a specific progenitor marker combination

To assess whether the putative differences in neurogenic capacity within the very same rhombomere could be foreshadowed by differential gene expression, we gene-profiled embryonic hindbrains during a temporal period encompassing the first and the second neurogenesis phases. To position the cells along the anteroposterior (AP) axis within the very same rhombomere, cells at the centers vs. cells at the neighboring regions, we resorted to zebrafish transgenic lines such as Mü4127 and Tg[elA:GFP], which label rhombomeres (r) 3 and 5 in DsRed and GFP, respectively (Distel et al., 2009; Labalette et al., 2011), and Tg[BCP:H2AmCherry/H2B-GFP] as landmarks of hindbrain boundaries (Hevia et al., 2022). First, we assessed the expression of *sox9b*, a transcription factor used as a marker of neural progenitors with radial glial features, which is necessary to maintain cells in the neural progenitor state but not sufficient for the generation of glial cells (Stolt et al., 2003). From 30hpf to 36hpf, *sox9b* expression was enriched in rhombomere centers (Figure 1A—B). After 36hpf, *sox9b* became homogenously expressed along the AP axis (Figure 1C). During this period, rhombomere centers displayed other well-described genes such as *meteorin*, *meteorin-like* as well as *erm,* the read-out of FGF-activity (Figure 1D—F). Next, we profiled hindbrains with markers such *fabp7a* and *slc1a2a*, which expression features radial glia progenitors during embryogenesis (Hartfuss et al., 2001; Than-Trong and Bally-Cuif, 2015). The expression of these specific radial glia markers revealed a delay in their onset of expression in the rhombomere centers with respect to *sox9b*. Expression of *fabp7a* was first observed at 30hpf, restricted to discrete groups of cells located mediolaterally all along the AP (Figure 1G). Later, *fabp7a* expression became enriched within stripes (Figure 1H—J), which corresponded to rhombomere centers (Figure K—K’), until at least 54hpf. Similarly, the onset of *slc1a2a* expression was between 30 and 39hpf (Figure 1L—M), and the enrichment was sustained until 54hpf (Figure 1M—O), a period when most of the cells within the adjacent regions had already engaged in neurogenesis (Gonzalez-Quevedo et al., 2010). Further, we confirmed that both *fabp7a* and *slc1a2a* were expressed in the same cells within the rhombomere centers (Figure 1P—P’). Overall, the spatiotemporal analysis of progenitor cell markers within the rhombomeres reveals that rhombomere centers show a specific combinatory of gene expression, with the enrichment of some radial glia markers.

**Figure 1:**
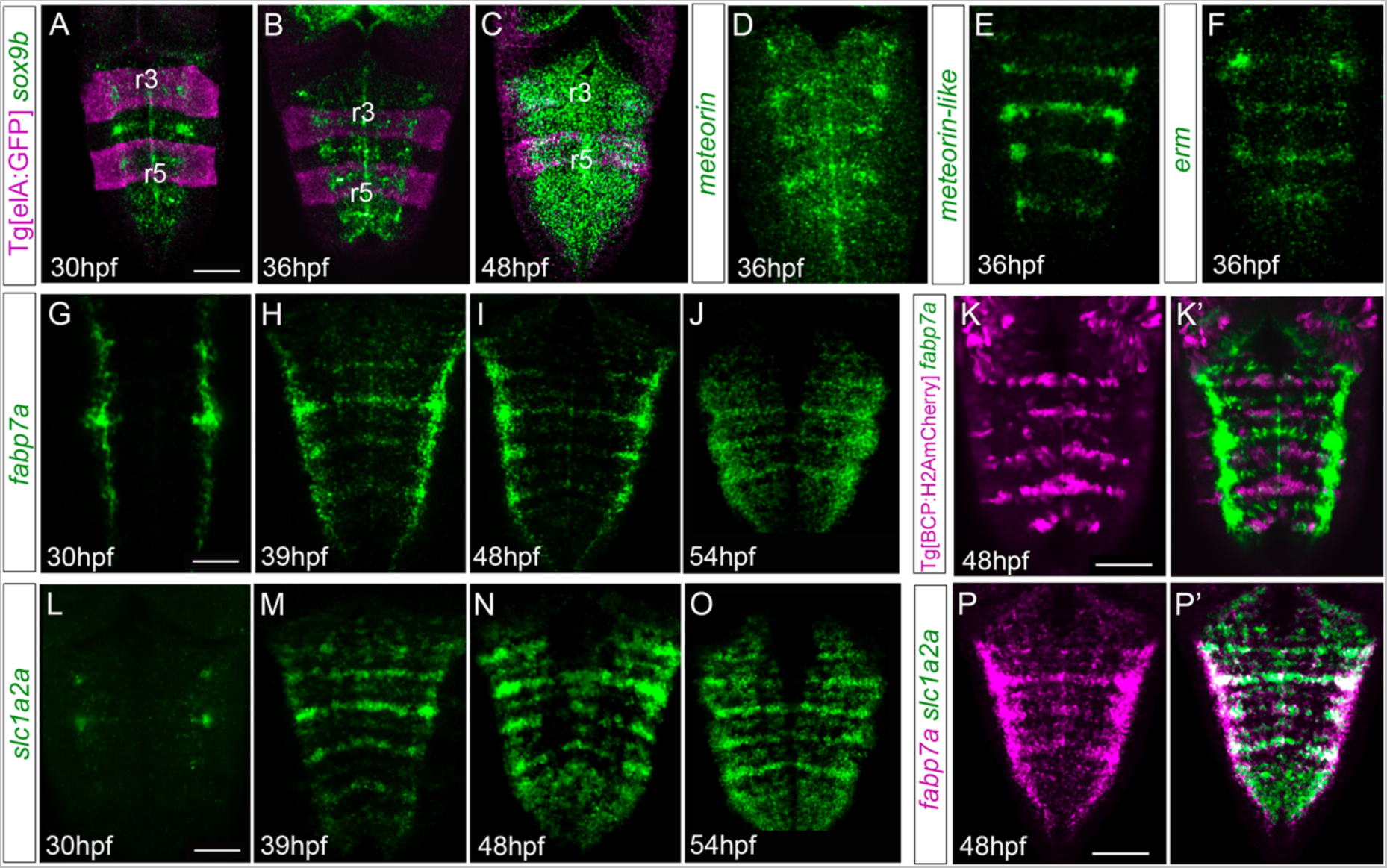
Hindbrain rhombomere centers display a specific combinatory of gene expression. (A—C) Tg[elA:GFP], (D—J, L—P, P’) wild type, or (K, K’) Tg[BCP:H2AmCherry] embryos at the indicated embryonic stages were *in situ* hybridized with *sox9* (A—C), *meteorin* (D), *meteorin-like* (E), *erm* (F), *fabp7a* (G—K, K’, P, P’), and *slc1a2a* (L—P, P’). Images are displayed as the single gene expression channel (D—K, L—P) or the overlay of the two corresponding channels (A—C, K’, P’). All images are dorsal Maximal Intensity Projections (MIP) with anterior to the top. Note the enrichment of these markers in rhombomere centers. r3 and r5, rhombomeres 3 and 5. Scale bar, 50μm.

### The centers of the rhombomeres harbor non-committed and actively proliferating progenitor cells

Next, we sought to investigate the type of progenitors present in the rhombomere centers. We first analyzed the expression of neural markers such as nestin and *sox2*, which are shared between neuroepithelial cells and radial glia progenitors. We observed that nestin was expressed in the hindbrain ventricular zone, without any spatial restriction along the AP axis (Figure 2A, a). Accordingly, *sox2* –another pan-neural progenitor marker– was expressed in the ventricular domain all along the AP axis and did not overlap with HuC, a pan-neuronal differentiation marker expressed in the neuronal differentiation domain (Figure 2B, b). To map the asynchronous patterning of neurogenesis within individual rhombomeres, we performed a colocalization analysis of *fabp7a* and the proneural genes *neurog1* and *ascl1b*, which label cells committed to the neuronal lineage (Bertrand et al., 2002). Double *in situ* hybridization experiments revealed that cells in the center of the rhombomeres (*fabp7a* cells) did not display *neurog1* and *ascl1b* expression, which were restricted to the neighboring domains (Figure 2C—D), the boundary-flanking regions (Nikolaou et al., 2009), and in a more ventral cell layer corresponding to the neurogenic committed regions (Belzunce et al., 2020; Hevia et al., 2022). Accordingly, *fabp7a* was expressed in the most ventricular progenitor cells (Figure 2c—d). These observations indicate that at this stage (48hpf) cells within rhombomere centers are non-committed progenitors, whereas progenitors in the boundary-flanking regions had already engaged in neurogenesis (Gonzalez-Quevedo et al., 2010).

**Figure 2:**
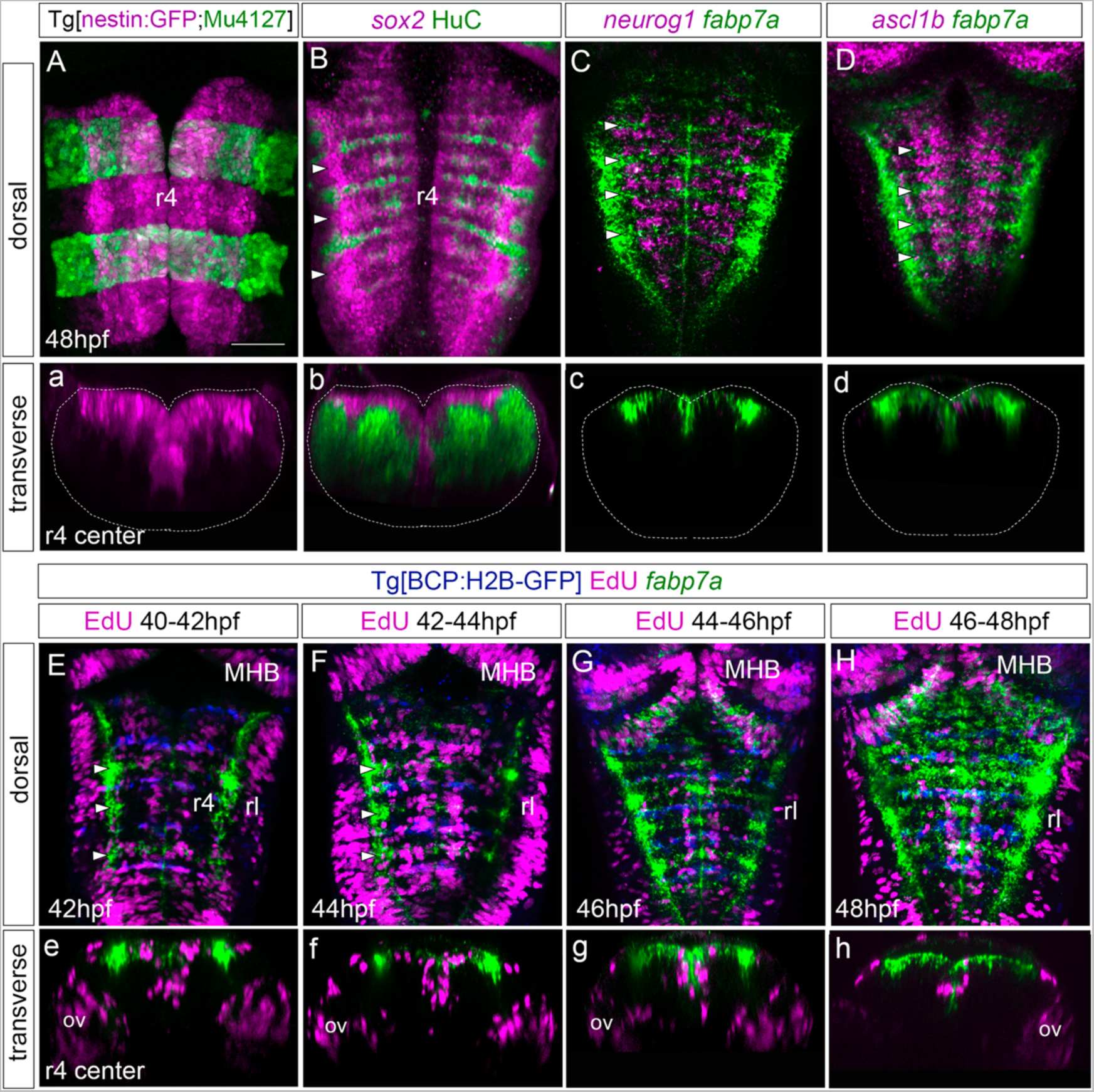
Rhombomere centers harbor proliferating neural progenitors. (A) Tg[nestin:GFP;Mu4127] embryos at 48hpf displaying neural progenitors in magenta and the r3 and r5 landmark in green. (B) Wild type embryos immunostained with anti-Sox2 (magenta) to visualize the neural progenitors and with anti-HuC (green) to label the differentiated neurons at 48hpf. (C—D) Wild type embryos *in situ* hybridized with *fabp7a* (green) to stain progenitors and *neurog1* or *ascl1b* (magenta) to label neuronal committed cells at 48hpf. (a—d) Transverse views of (A—D), through the center of r4. (a) Images displaying a single channel or (b—d) the overlay of both channels. (E—H) Tg[BCP:H2B-GFP] embryos were incubated with EdU for 2h, *in situ* hybridized with *fabp7a* and immunostained with anti-GFP to label boundary cells. The EdU-positive cells are displayed in magenta, the position the boundaries in blue, and *fabp7a* expression indicating the center of the rhombomeres in green. (e—h) Transverse views of (E—H), through r4 center displaying only the magenta and green channels. (A—H) Dorsal MIP with anterior to the top displaying all channels. White arrowheads indicate the center of rhombomeres. Dotted lines in (a—d) indicate the contour of the neural tube. ov, otic vesicle; rl, rhombic lip. Scale bar, 50μm.

With the increasing evidence that cells within rhombomere centers display distinct features than the neighboring regions, we wanted to seek whether these molecular differences translated into distinct proliferative behaviors. First, we studied the putative changes on proliferative activity by EdU-incorporation experiments. To reveal which cells underwent DNA replication, we shortly pulsed Tg[BCP:H2B-GFP] embryos displaying GFP in the hindbrain boundaries (Hevia et al., 2022), with EdU at different time intervals. This was followed by *fabp7a* staining to label the cells in the rhombomere centers (Figure 2E—H; e—h). We observed that during 40-42hpf and 42-44hpf intervals many cells in the whole the hindbrain incorporated EdU, including the mid-hindbrain boundary (MHB) and the rhombic lip (rl) (Figure 2E—F), suggesting that cells at the center of the rhombomere underwent S-phase during these periods although not all *fabp7a* cells incorporated EdU (Figure 2e—f). However, despite in the 44-46hpf EdU-pulse period, many *fabp7a*-cells were still proliferating (Figure 2G, g), the proliferative capacity of the overall hindbrain progenitors diminished. And by 46-48hpf almost no cells within the rhombomere centers had incorporated EdU (Figure 2H, h), except for some EdU-positive cells in the more medioventral domain were found (Figure 2g— h), suggesting that the derivatives of rhombomere centers might be neurons. These results indicate that the centers of the rhombomeres harbor proliferating progenitors that seem to diminish their proliferative capacity from 46hpf onwards.

### The progenitor cells within the rhombomeric centers are maintained in the G1 phase

Next, we explored whether progenitor cells that decreased the proliferative capacity were arrested in a specific cell cycle phase. We performed *in vivo* analysis of PCNA-GFP dynamics by injecting it in Mu4127 embryos, expressing DsRed in rhombomeres 3 and 5. We imaged PCNA-GFP before and after these cells decreased the proliferative activity, at 42hpf and 48hpf, and focused on the PCNA-GFP localization patterns, particularly on cell nuclei displaying a finely speckled signal as a mark of S-phase (Figure 3A). Quantitative analysis at 42hpf showed that both rhombomere centers and the adjacent flanking regions contained a similar percentage of cells in S-phase (Figure 3B; 8.2% in rhombomeres centers vs. 10.5% in flanking regions). However, at 48hpf, the S-phase cells in the rhombomere centers decreased significantly (Figure 3C; 3.2% in rhombomeres centers vs. 9.1% in flanking regions). This evidence let us to explore further the possibility that these progenitors could be arrested in the cell cycle. For this, we made use of the Tg[Fucci] line (Bouldin and Kimelman, 2014), in which cells in the G1-phase display mCherry-zCdt1. We pulsed embryos with EdU at 46hpf for 2h —the time proliferation started to decrease— and image them to assess the colocalization of EdU-positive cells (S-phase cells) and fucci cells (G1-phase cells) within the rhombomeres. G1-phase cells were enriched in the center of the rhombomeres, with a very similar pattern to *fabp7a* (Figure 3D—D’) and, as expected, EdU-positive cells did not display the G1-phase marker (Figure 3d—d’). These evidences suggest that the rhombomere centers from 46hpf onwards harbor cells that may remain as quiescent progenitors.

**Figure 3:**
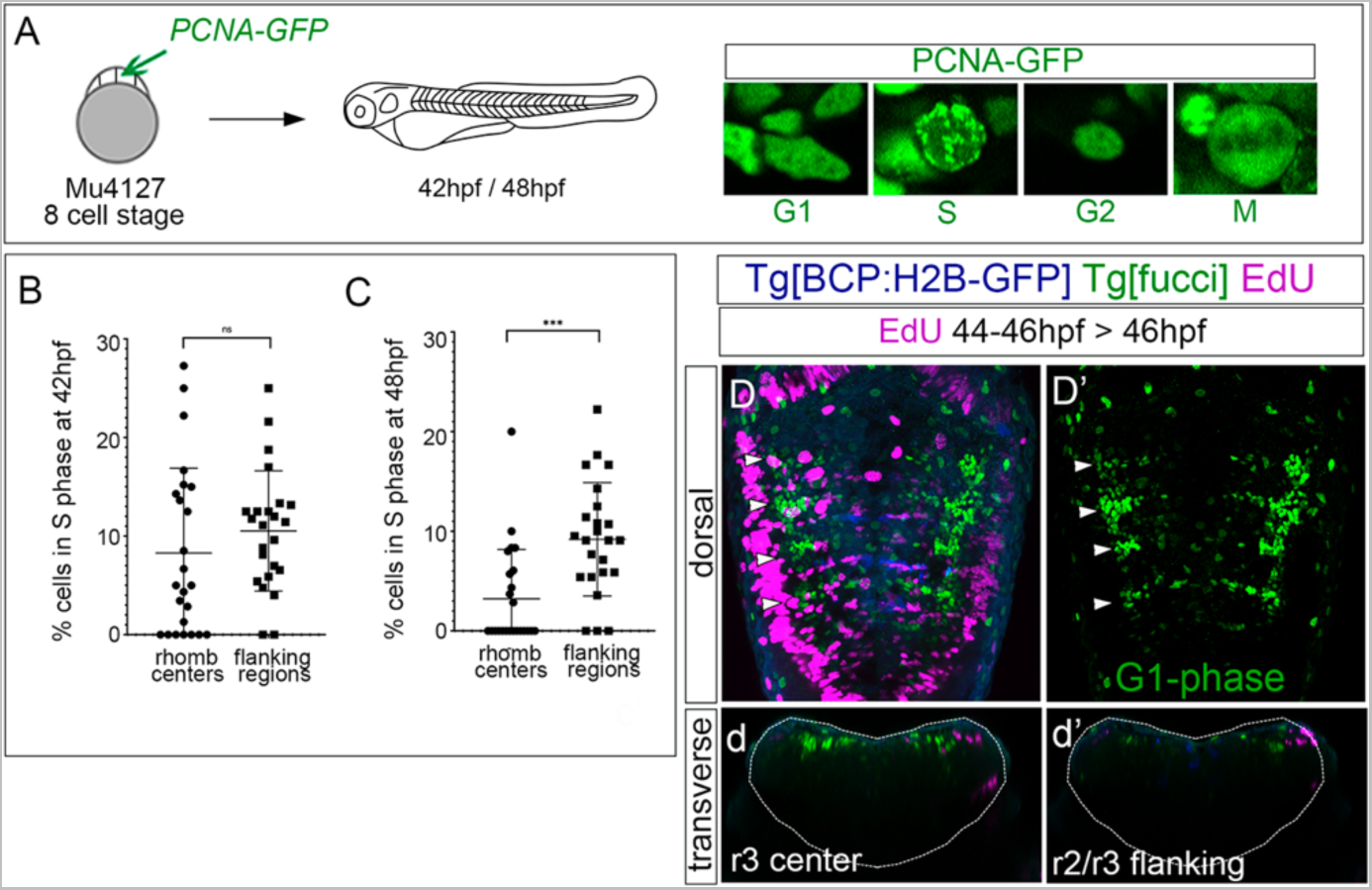
The rhombomere centers harbor G1-phase arrested progenitors. (A) Scheme depicting the experimental design of the *in vivo* PCNA-GFP clonal analysis. Mu4127 embryos, displaying DsRed in r3 and r5, were injected with PCNA-GFP at 8-cell stage and hindbrains were analysed at 42hpf and 48hpf. Images depict examples of the different distribution of PCNA-GFP within the cell nuclei along the the cell cycle phases. (B—C) Graphs illustrating the percentage of cells in S-phase in rhombomere centers and boundary flanking regions at 42hpf and 48hpf, respectively. S-phase cells at 42hpf: 8.2% in rhombomere centers vs. 10.5% in boundary flanking regions. S-phase cells at 48hpf: 3.2% in rhombomere centers vs. 9.1% in boundary flanking regions. Wilcoxon test analysis are shown. ns, non-significant, ***p<0.001, N = 4 embryos, n = 24 boundaries, n = 24 flanking regions. (D—D’) Double transgenic Tg[BCP:H2B-GFP;fucci] embryos were incubated with EdU for 2 hours and analyzed at 46hpf. Boundaries are depicted in blue, G1-phase cells in green and EdU-positive cells in magenta. Note that G1-phase cells are mainly located in the center of the rhombomeres and did not incorporate EdU. (D) displays a merge of channels and (D’) only the green channel (G1-phase cells). (d—d’) Transverse views of (D) at the level of r3 and r2/r3 flanking regions, respectively. (D—D’) Dorsal MIP with anterior to the top. Dotted lines in (d— d’) indicate the contour of the neural tube.

### Proliferative progenitors in rhombomere centers exhibit different division modes

Next, to assess the derivatives of the progenitors at the rhombomere centers, we performed EdU-pulse-and-chase experiments, which allows us to analyze the final position of the cells that incorporated EdU at the given pulse time. Thus, Tg[BCP:H2B-GFP] embryos after a 2h EdU-pulse were chased for 6h and *in situ* hybridized with *fabp7a.* Then, the position of EdU-positive cells in the ventricular vs. neuronal differentiated domain was assessed (Figure 4A— D). Although at 48hpf, most of the EdU-positive cells remained in the ventricular progenitor zone and expressed *fabp7a*, some were found in the neuronal differentiation domain (Figure 4a), indicating that most probably they underwent neuronal differentiation. In line with our previous results, the *fabp7a* cells in S-phase diminished over time (Figure 4B—C, b—d), with a dramatic decrease of cells that incorporated EdU from 46hpf onwards (Figure 4D, d). Despite our results suggest that progenitor cells within rhombomere centers can give rise to neuronal derivatives, we cannot rule out whether the remaining progenitors in the ventricular zone are slow-dividing progenitors holding longer cell cycles that last more than 8h, described as the estimated average time for hindbrain cells to undergo división (Lyons et al., 2003), or whether they are kept in quiescence arrested out of the cell cycle.

**Figure 4:**
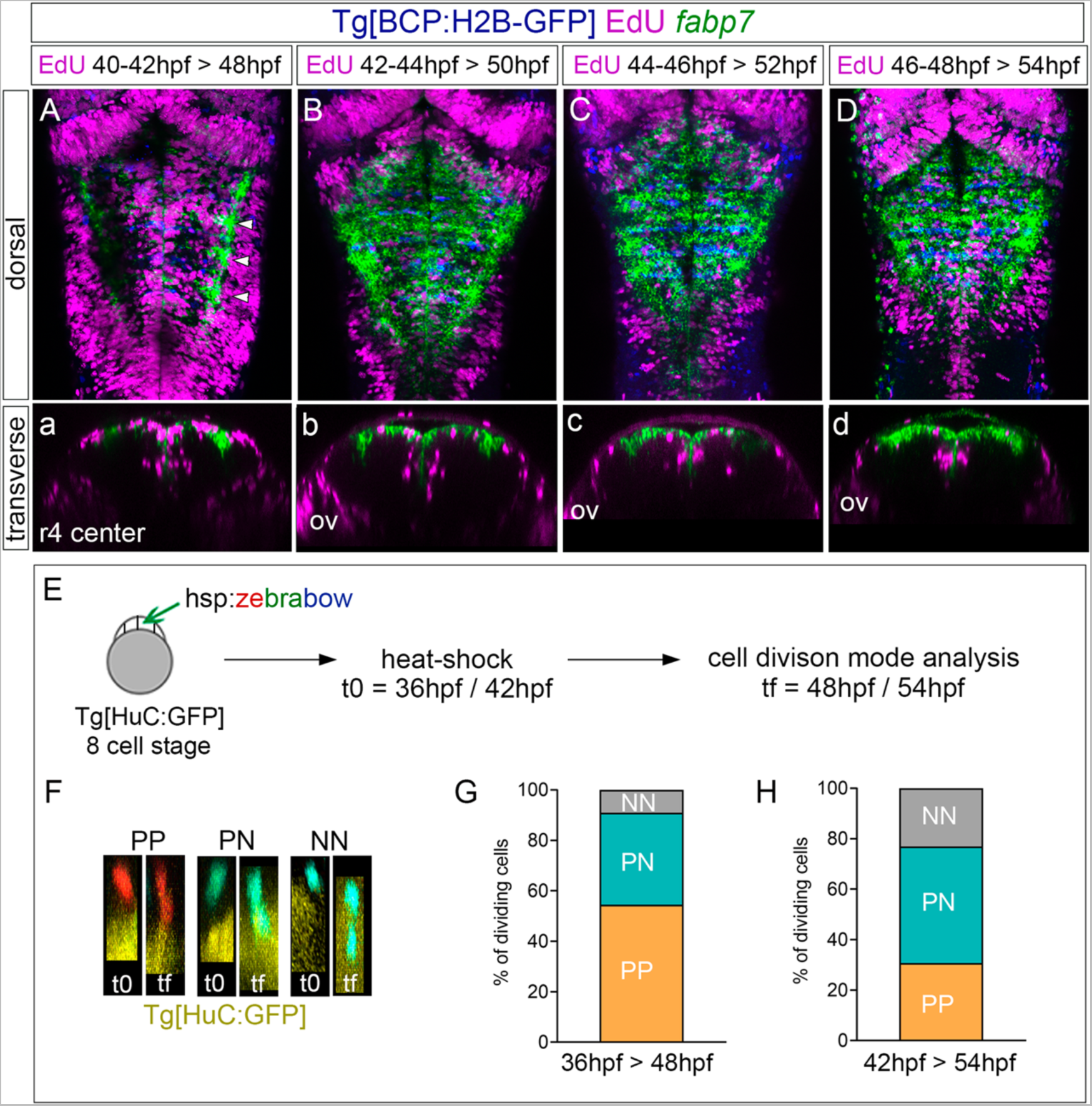
The proliferating progenitor cells within the rhombomere centers shift their division mode over time. (A—D) Tg[BCP:H2B-GFP] embryos were pulsed with EdU at the indicated intervals, and chased for 6 hours, to analyze the position of the derivatives of cells that underwent S-phase. EdU was detected in magenta, hindbrain boundaries in blue, and *fabp7a* cells marking the center of the rhombomeres in green. (A—D) Dorsal projections with anterior to the top. (a—d) Transverse views of (A—D) through the center of r4, displaying only the magenta and green channels. ov, otic vesicle. (E) Analysis of the progenitor cell division mode within the rhombomere centers. Scheme depicting the multicolor clonal growth experimental approach. Tg[HuC:GFP] embryos at 8-cell stage were injected with the hsp:zebrabow construct and heat shocked 3 hours before the imaging time. Embryos were *in vivo* imaged at 36hpf or 42hpf (t0), and at 48hpf or 54hpf (tf), respectively, and the cell division mode was analyzed in all the clones. (F) Transverse projections generated at t0 and tf showing examples of the three observed clonal cell behaviors: a cell undergoing symmetric proliferative division (PP, red cell not expressing HuC at t0 and the daughter cells not expressing it at tf), a cell undergoing asymmetric division (PN, turquoise cell not expressing HuC at t0, with only one of the daughters expressing HuC at tf), and a cell undergoing symmetric neurogenic division (NN, turquoise cell not expressing HuC at t0, but both daughters expressing HuC at tf). The neuronal differentiation domain is displayed in yellow. (G—H) Stacked bar graphs showing the percentage of PP (orange), PN (blue) and NN (grey) cell division modes in the rhombomere centers between two different time intervals: 36-48hpf (G, n= 11), and 42-54hpf (H, n = 13).

Since progenitor cells within the rhombomere centers seem to contribute to the neuronal lineage, the next step was to unveil their cell division mode. For this, we used the zebrabow multicolor clonal analysis approach (Brockway et al., 2019), which allows us to label in the same color all the derivatives from a given cell clone. We classified the different division modes according to two criteria: the relative position between sister cells after division and the expression of the differentiation marker HuC. Tg[HuC:GFP] embryos at 8 cell-stage were injected with the hsp:zebrabow construct, heat-shocked at two different times (36hpf or 42hpf), and let to develop for 6 more hours (Figure 4E). Then, the percentage of cells in the rhombomere centers undergoing the different division modes was assessed: symmetric proliferative, giving rise to two progenitor cells (PP); asymmetric, giving rise to one progenitor and one neuronal derivative (PN); and symmetric neurogenic, giving rise to two neurons (NN) (Figure 4F). Although at 48hpf most cell divisions were symmetric proliferative (Figure 4G; PP: 54.5% vs. PN: 36.3% vs. NN: 9.2%), by 54hpf the asymmetric and neurogenic divisions increased at the expense of the proliferative divisions (Figure 4H; PP: 30.7% vs. PN: 46.2% vs. NN: 23.1%), supporting the change in behavior of these cells that we previously envisaged.

### Progenitor cells in the rhombomeric centers display Notch-activity

We were eager to understand by which mechanism cells within the center of the rhombomeres behave differently than their adjacent neighbors. Since we previously showed that Notch-signaling was important in governing the regulation of binary cell choices in the hindbrain boundaries (Hevia et al., 2022), we investigated whether Notch-activity played any role in the rhombomere centers. First, we assessed whether Notch-activity was spatiotemporally restricted within the hindbrain at these late embryonic stages using a readout of Notch-active cells, the Tg[tp1:d2GP] line (Clark et al., 2012). At 24hpf, rhombomeres behave as proneural clusters (Nikolaou et al., 2009), and therefore Notch-activity is distributed in the whole segment (Hevia et al., 2022). However, by 48hpf there was an enrichment of Notch-activity within the center of the rhombomeres (Figure 5A) together with the hindbrain boundaries (Hevia et al., 2022). Notch-activity remained in the rhombomere centers at least until 54hpf (Figure 5B) and it was confined to progenitor cells displaying radial projections towards the mantle zone (Figure 5a—b). To demonstrate that *fabp7a* cells were Notch-active, we labelled Tg[tp1:d2GP] embryos with fabp7a and revealed that indeed most of the fabp7a-cells in the rhombomere centers displayed Notch activity (Figure 5C, c—c’’). These results indicated that progenitor cells within the rhombomeric centers are Notch-active concomitantly with the enrichment of *fapb7a* and *slc1a2a* expression in these domains. Next, we analyzed the expression of Notch signaling players within the hindbrain. First, we studied the expression of Notch receptors focusing on *notch3*, since it was described to be crucial in maintaining progenitor cells within a non-committed progenitor state (Than-Trong et al., 2018; Hevia et al., 2022). The spatiotemporal analysis of *notch3* revealed its faint expression in the hindbrain progenitor domain by 30hpf (Figure 5D; (Hevia et al., 2022), although it increased by 36hpf and was maintained at least until 54hpf (Figure 5E—F). Colocalization analysis at 48hpf revealed the overlapping expression of *notch3* and *fabp7a* in the rhombomere centers (Figure 5G, g—g’’), supporting the idea that *notch3* might maintain this cell population in a non-committed progenitor state. However, both *notch1a* and *notch1b* were as well expressed in the rhombomeric centers and overlapped with *fabp7a* (Supplementary Figure 1). To further explore the complex Notch network within this cell population, we investigated the expression of the Notch-signaling targets. Previous work in adult zebrafish pallium described that quiescent radial glia cells expressing *fabp7a* maintain their stemness features upon Notch3 signaling by the target gene *hey1* (Than-Trong et al., 2018). Moreover, in the zebrafish retina *hey1* regulates the Müller glia downstream *notch3* thereby reducing their proliferative capacity upon injury (Sahu et al., 2021). Thus, these evidences drove us to *hey1* as a good candidate downstream of *notch3*. We performed the spatiotemporal expression analysis of *hey1* within the zebrafish hindbrain. Although at 24hpf, no expression of *hey1* was observed in the hindbrain (result not shown), by 36hpf *hey1* was specifically expressed in rhombomere centers and remained at least until 54hpf (Figure 5H—J). *hey1* overlapped with *fabp7a* at the center of the rhombomeres (Figure 5K), although not all *fabp7a* progenitors displayed *hey1* (Figure 5k—k’’), suggesting that Notch-pathway may operate through *hey1* in subpopulation in rhombomere centers.

**Figure 5:**
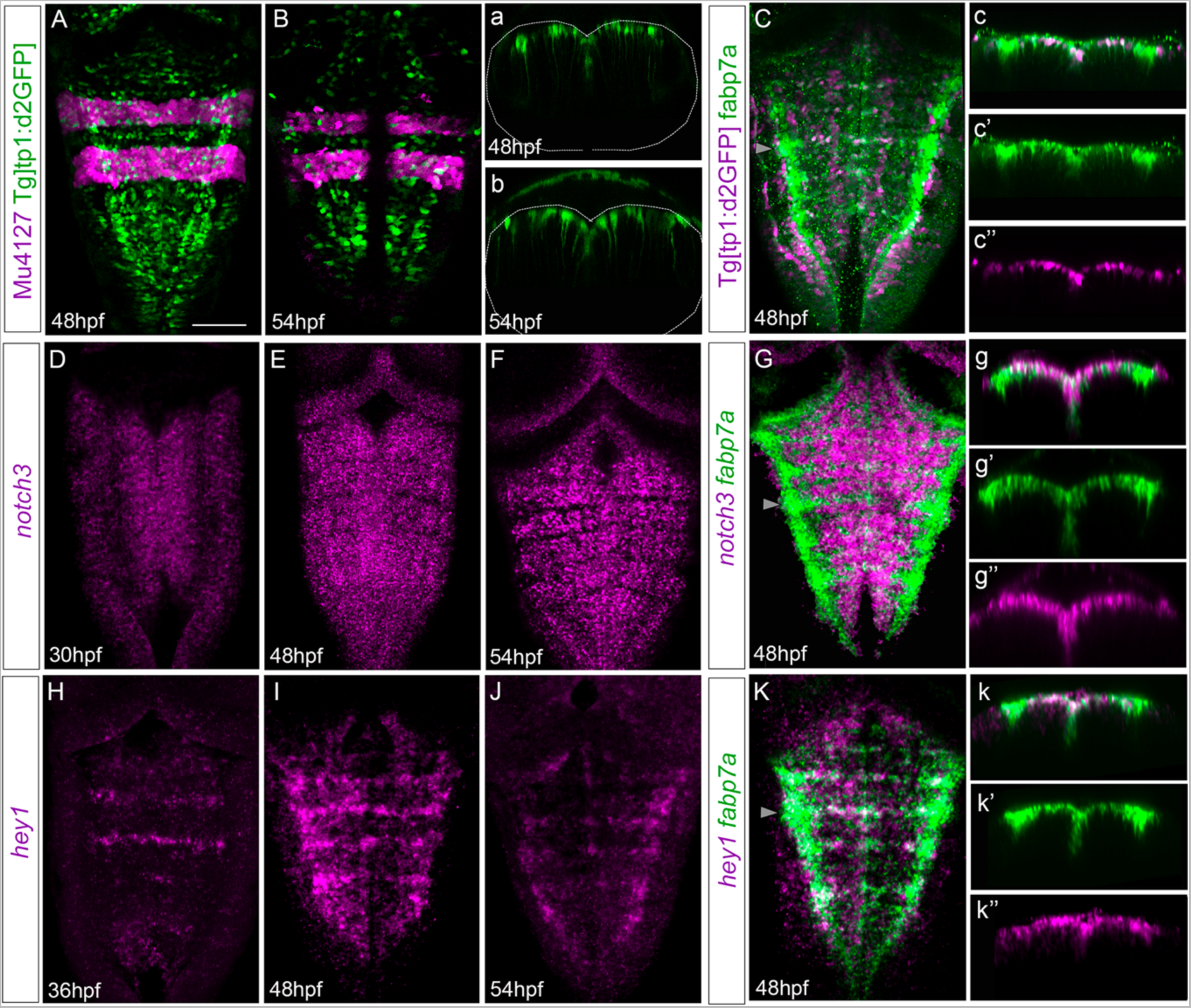
Rhombomere centers display Notch-activity. (A—B) Tg[tp1:d2GFP;Mu4127] embryos as readout of Notch-activity (green) with the r3 and r5 landmarks (magenta) at the indicated stages. (a—b) Transverse view of (A—B), through the center of r4 displaying only the Notch-active cells. Dotted lines indicate the contour of the neural tube. (C) Tg[tp1:d2GFP] embryos displaying Notch activity (magenta) and immunostained with fabp7a (green) at 48hpf. (c—c’’) Transverse view of (C) showing only the ventricular domain through the center of r4 (grey arrowhead) displaying either both channels (c), or single channels (c’—c’’). (D—G) Wild-type embryos *in situ* hybridized either with *notch3* (D—F), or *notch3* and *fabp7a* (G) at the indicated stages. (g—g’’) Transverse view of (G) showing only the ventricular domain through the center of r4 (arrowhead) displaying either both channels (g), or single channels (g’—g’’). (H—K) Wild-type embryos were *in situ* hybridized either with *hey1* (H—J), or *hey1* and *fabp7a* (K) at the indicated stages. (k—k’’) Transverse view of (K) showing only the ventricular domain through the center of r4 (arrowhead) displaying either both channels (k), or *fabp7a* or *hey1* (k’—k’’). (A—K) Dorsal MIP with anterior to the top. Scale bar, 50μm.

### Notch3 is necessary for the maintenance of cells within rhombomere centers as non-committed progenitors

The display of Notch-activity and the expression of Notch players within rhombomere centers led us to seek whether these progenitors responded to Notch. For this, we conditionally abrogated Notch signaling using the pharmacological reagent LY411575, which is a gamma-secretase inhibitor resulting in the downregulation of the Notch pathway. First, we inhibited Notch-activity by incubating Tg[tp1:dGFP] embryos with either DMSO as control (Figure 6A— E) or LY411575 (Figure 6F-J) from 36 to 42hpf (Figure 6A—J), before rhombomere centers decrease their proliferative capacity, and analyzed the effects on progenitor cells, committed progenitors, differentiated neurons and on the Notch target *hey1*. Upon abrogation of Notch activity, expression of *fabp7a* and *slc1a2a* progenitor markers was completely downregulated when compared to control embryos (Figure 6A—B, F—G). Neurogenesis and neuronal differentiation increased upon Notch-inhibition and the well described proneural genes salt-and-pepper expression pattern was lost (Figure 6C, H), demonstrating that cells dramatically engaged in neurogenesis. Accordingly, there was an increase of the neuronal differentiation domain at expense of the progenitor domain (Figure 6D, I). Moreover, the expression of the Notch-target *hey1* was completely inhibited (Figure 6E, J), suggesting that Notch-activity was acting through *hey1* in the rhombomere centers. These results indicated that Notch signaling maintains hindbrain progenitor cells –including those within rhombomere centers– as non-committed progenitors. To dissect whether Notch-pathway was important for maintaining these cells in the progenitor state at later stages, we inhibited Notch in a later temporal window, from 48 to 54hpf, and observed a similar phenotype (Figure 6K—N). When we quantified the volume of the progenitor vs. the neuronal differentiation domains in control and LY41575-treated embryos at different stages we confirmed this phenotype (Supplementary Figure 2A—B), and demonstrated it was not a result of an increase of cell death (Supplementary Figure 2C).

**Figure 6:**
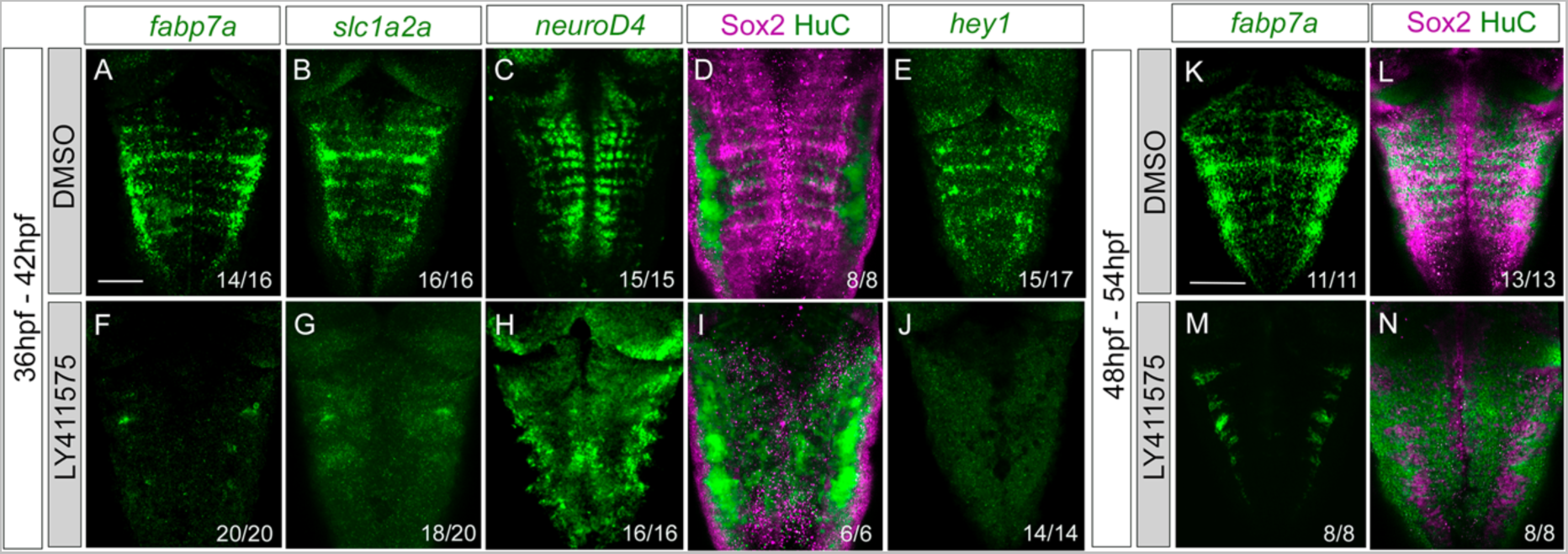
Progenitor cells in the rhombomere centers are Notch-responsive. Wild type embryos were treated with either DMSO (A—E, K—L) or the gamma secretase inhibitor LY411575 (F—J, M—N) for 6 hours, at 36hpf (A—J) or at 48hpf (K—N). Embryos were *in situ* hybridized with *fabp7a* (A, F, K, M), *slc1a2a* (B, G), *neuroD4* (C, H), and *hey1* (E, J), or immunostaining with Sox2 and HuC (D, I, L, N). Dorsal MIP with anterior to the top. Numbers at the bottom indicate the individuals with the displayed phenotype over the total of analyzed specimens. Scale bar, 50μm.

To study the contribution of the Notch3-signaling to the Notch-activity we inhibited the expression of *notch3* by the use of the null *notch3^fh332/fh332^* mutants (Alunni et al., 2013) combined with the Notch activity reporter. Tg[tp1:d2GFP;*notch3^fh332/fh332^*] embryos analysis revealed a dramatic decrease of the patterned Notch-activity in the hindbrain (Figure 7A, F). Next, we analyzed whether Notch3 was involved in ascribing the progenitor capacities to this specific cell population. *notch3* mutation resulted in a loss of *fabp7a* expression in the rhombomere centers (Figure 7B, G), demonstrating the relevant contribution of Notch3-activity in the *fabp7a* progenitors. Expression of the proneural genes *ascl1b* and *neuroD4* was affected as well (Figure 7C—D, H—I), as a consequence of all progenitors undergoing neuronal differentiation. The expression of the Notch target *hey1* was also downregulated (Figure 7E— J). To account for any role for apoptosis within this phenotype, we analyzed cell death figures and no differences in the number of apoptotic events were observed between wild type and *notch3^fh332/fh332^* mutants (wild type: 0.4 ± 0.8 apoptotic cells n=16 vs. *notch3^fh332/fh332^* 1 ± 2.2 apoptotic cells n=14, ns). Therefore, these results indicate that *notch3* is necessary for the maintenance of the progenitor state within this cell population. However, *fabp7a* and proneural gene expression analyses in *hey1^ha11/ha11^* mutants did not show any differences when compared with control embryos (K—N), suggesting that *hey1* might not be the Notch3 effector in maintaining the *fabp7a* cells as progenitors during this time window.

**Figure 7:**
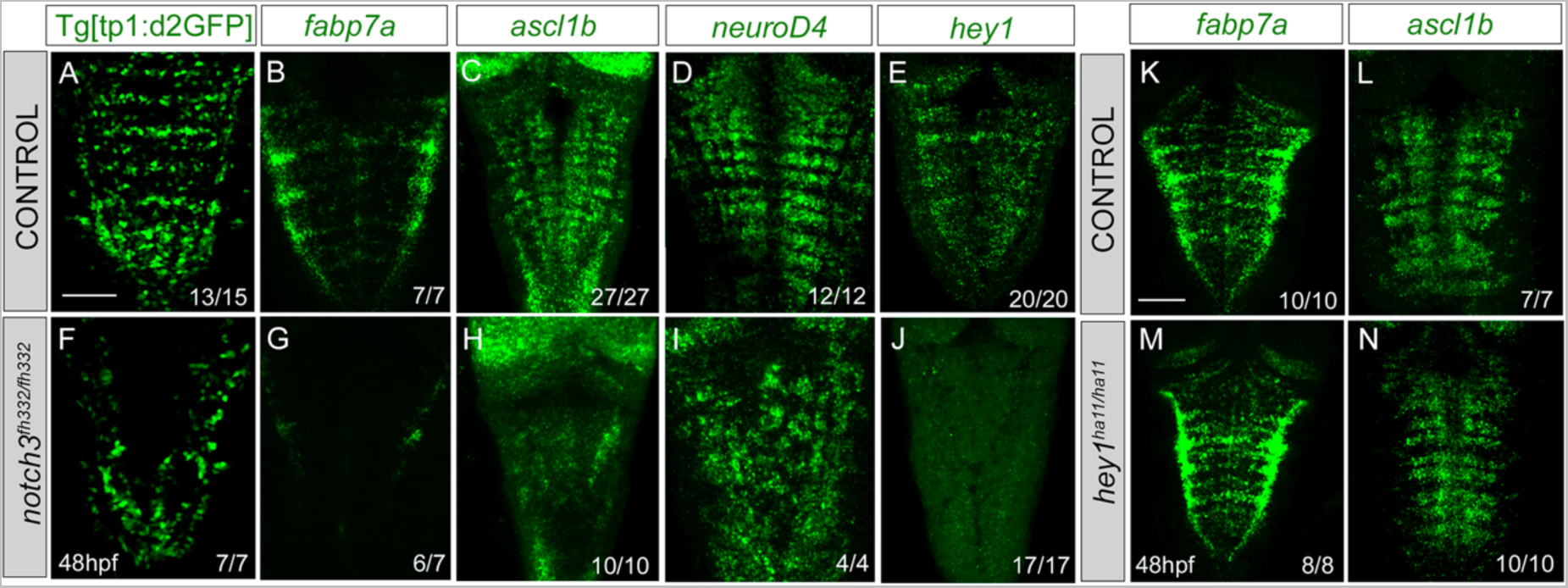
***notch3* mutation accounts for the loss of Notch-activity in the center of the rhombomeres and results in neuronal differentiation.** (A—J) Control and *notch3^fh322/fh322^* embryos in the Tg[tp1:d2GFP] background were *in situ* hybridized with *fabp7a* (B, G), *ascl1b* (C, H), *neuroD4* (D, I), or *hey1* (E, J) at 48hpf. (K—N) Control and *hey1^ha11/ha11^* embryos were in *in situ* hybridized with *fabp7a* (K, M), and *ascl1b* (L, N) at 48hpf. Dorsal MIP with anterior to the top. Numbers at the bottom indicate the individuals with the displayed phenotype over the total of analyzed specimens. As control embryos, either wild type or heterozygous were used. Scale bar, 50μm.

## DISCUSSION

Previous studies have described that differential neurogenesis in the hindbrain is spatiotemporally regulated. During the first neurogenesis phase, rhombomeres act as proneural clusters (Nikolaou et al., 2009), whereas hindbrain boundaries are devoid of neurogenesis (Voltes et al., 2019; Hevia et al., 2022). However, later on, boundary progenitor cells transition toward neurons relying on Notch3-signaling (Hevia et al., 2022). This distinct neurogenic behavior is usually foreshadowed by different gene expression, since boundaries express a specific gene combination (Cheng et al., 2004; Letelier et al., 2018). Interestingly, differences in the neurogenic capacity are generated within the same rhombomere, since neurogenesis gets restricted to the boundary flanking regions whereas the centers of the rhombomeres instruct the position of the neighboring neurons through FGF-signaling (Gonzalez-Quevedo et al., 2010). This confinement of the neurogenesis to the boundary-flanking regions is regulated by miR-9, which exerts distinct actions at different stages of progenitor commitment along the neurogenesis cascade (Coolen et al., 2012). Thus, the center of the rhombomeres work as a signaling hub, that later on will express radial glia cell markers and will provide glial cells (Esain et al., 2010). Our results provide evidences to foresee another layer of complexity in the attribution of different progenitor behaviors, and stress the importance of the spatiotemporal coordination of neurogenesis. The enrichment in *sox9, meteorin*, and *meteorin-like* in the rhombomere centers during a short period of time, just before the onset of radial glia cell markers, could keep this cell population out of the early neurogenic program the neighboring cells already engaged in. This occurs at the same time that FGF signaling from neurons in the ventral hindbrain is required to downregulate proneural gene expression (Gonzalez-Quevedo et al., 2010). Thus, it seems that FGF-signaling represses proneural genes at the centers permitting the *fabp7a*-cells to proliferate.

One strategy to properly organize the generation of the neuronal circuits is to restrict the progenitor capacities within specific territories and to maintain groups of long-lasting progenitors (for review see (Belmonte-Mateos and Pujades, 2022). Several examples of the existence of distinct progenitor pools during embryogenesis (for review see (Stigloher et al., 2008), and the maintenance of quiescent cells in adult organisms (Than-Trong and Bally-Cuif, 2015; Than-Trong et al., 2018; Urbán et al., 2019) have been described. Our data show that this could also occur in the hindbrain. We demonstrate for the first time the center of the rhombomeres harbor proliferating progenitors since embryos display EdU labeling across the hindbrain including the rhombomeric centers. However, at a later temporal window, beyond 48hpf, their division capacity decreases significantly and fewer cells are in S-phase. Accordingly, we could demonstrate that beyond 46hpf the centers of the rhombomeres are enriched in progenitors in G1-phase, which considering the significant decrease in proliferation at these stages would indicate rhombomere centers harbor progenitors in a quiescent or slow division mode. These would suggest that different strategies are used within the same tissue to differentially pattern neurogenesis, or in order words, to restrict the progenitor features. Moreover, our zebrabow clonal analysis experiments also unraveled that of those progenitors that divide, they did it in different modes. And in line with our previous results, these confirm that at a later developmental window asymmetric and symmetric neurogenic cell divisions increased at the expense of symmetric proliferative divisions. Our results depict a scenario similar to what has been found in the mouse brain, where authors demonstrated the presence of slow-dividing progenitors in the embryonic telencephalon being the precursors of quiescent neural stem cells in the adult brain (Furutachi et al., 2015; Harada et al., 2021). It is therefore possible that cells in rhombomere centers are spatiotemporally regulated in their proliferative and neurogenic capacity to ensure the maintenance of progenitors in order to provide neurons at a later temporal window to maintain the homeostasis of the hindbrain.

What is the molecular mechanism responsible for the neurogenic patterning within the rhombomeres? Despite FGF20 being involved in the downregulation of proneural genes in rhombomere centers (Gonzalez-Quevedo et al., 2010), we explored the possible role of Notch signaling in this cell population for its role in the maintenance of slow-dividing/quiescent cells in the zebrafish pallium (Than-Trong et al., 2018). Cells in rhombomeres centers are Notch active from 48hpf onwards, and Notch signaling maintains them as non-committed progenitors. Although other Notch receptors are expressed earlier, Notch3 onset of expression coincides with the expression of *sox9b*, *meteorin* and *meteorin-like* in the centers and co-expressing as well with its target *hey1*. Thus, it seems that Notch3 is the main Notch receptor responsible of Notch-activity during later neurogenesis in the hindbrain (Hevia et al., 2022). Interestingly, it was described that the sustained expression of the Notch3 target hey1 maintained neural progenitors in a quiescent and slow dividing mode in both adult zebrafish telencephalon as well as the subventricular zone in mice embryonic brain, respectively (Furutachi et al., 2015; Than-Trong et al., 2018; Harada et al., 2021). Despite our functional studies demonstrate that cells in rhombomere centers rely on Notch3 to keep as non-committed progenitors, they also suggest *hey1* might not be the target through which it exercises its function within hindbrain rhombomeres. Most probably, the scenario is a bit more complex and other *hey* or *her* genes could compensate for the effects of abrogating *hey1*, as a way of providing robustness to the system.

## ACKNOWLEDGEMENTS AND FUNDING SOURCES

We would like to thank Laia Subirana and Marta Linares for technical assistance and members of the lab for insights and critical discussions. We thank L Bally-Cuif for the Tg[*notch3^fh332^*] and Tg[*hey^ha11^*] mutants, D Kimelman for providing the Fucci construct and TA Weissman for the hsp:zebrabow plasmid. Confocal microscopy was performed at the Advanced Light Microscopy Unit at the CRG. This work was funded by grants BFU2015-67400-P, PGC2018-095663-B-I00 and PID2021-123261NB-I00 from Ministerio de Ciencia e Innovación (MICIN), Agencia Estatal de Investigación (AEI, DOI: 10.13039/501100011033) and Fondo Europeo de Desarrollo Regional (FEDER) to CP. The Department of Medicine and Life Sciences (UPF) is a Unidad de Excelencia María de Maeztu (CEX2018-000792-M) funded by the MICIN and the AEI. CBM was recipient of predoctoral fellowship from the MICIN (FPI, BES-2016-076664). CP is a recipient of ICREA Academia award (Generalitat de Catalunya).

## AUTHOR CONTRIBUTIONS

CBM, LM and CP contributed to the concept and design of experiments. CBM and LM performed and analyzed the experiments. CBM, LM and CP interpreted the results. CBM and CP wrote the manuscript.

## DECLARATION OF INTERESTS

The authors declare no competing interests.

## CONTRIBUTION TO THE FIELD STATEMENT

In this work we show that neurogenesis along hindbrain rhombomeres is not homogeneous. Combining gene expression studies and cell proliferation experiments with cell lineage analyses we demonstrate that the neurogenic potential evolves during development and rhombomeres contain progenitor cells displaying different behaviors. Rhombomere centers harbor a heterogeneous population of proliferating progenitors expressing a given combination of radial glia cell markers. Some of these progenitors are arrested in G1-phase, whereas the others undergo cell division modes that shift over time, contributing to the neuronal lineages while maintaining the progenitor pool. They do so in an asynchronous manner to the neighboring territories. In addition, our functional experiments demonstrate that Notch is one of the key mechanisms governing the maintenance of these cells as non-committed progenitors. Overall, this study enlightens how cell fate commitment occurs asynchronously over time across different progenitor populations during hindbrain morphogenesis

## SUPPLEMENTARY MATERIAL

**Supplementary Figure 1:**
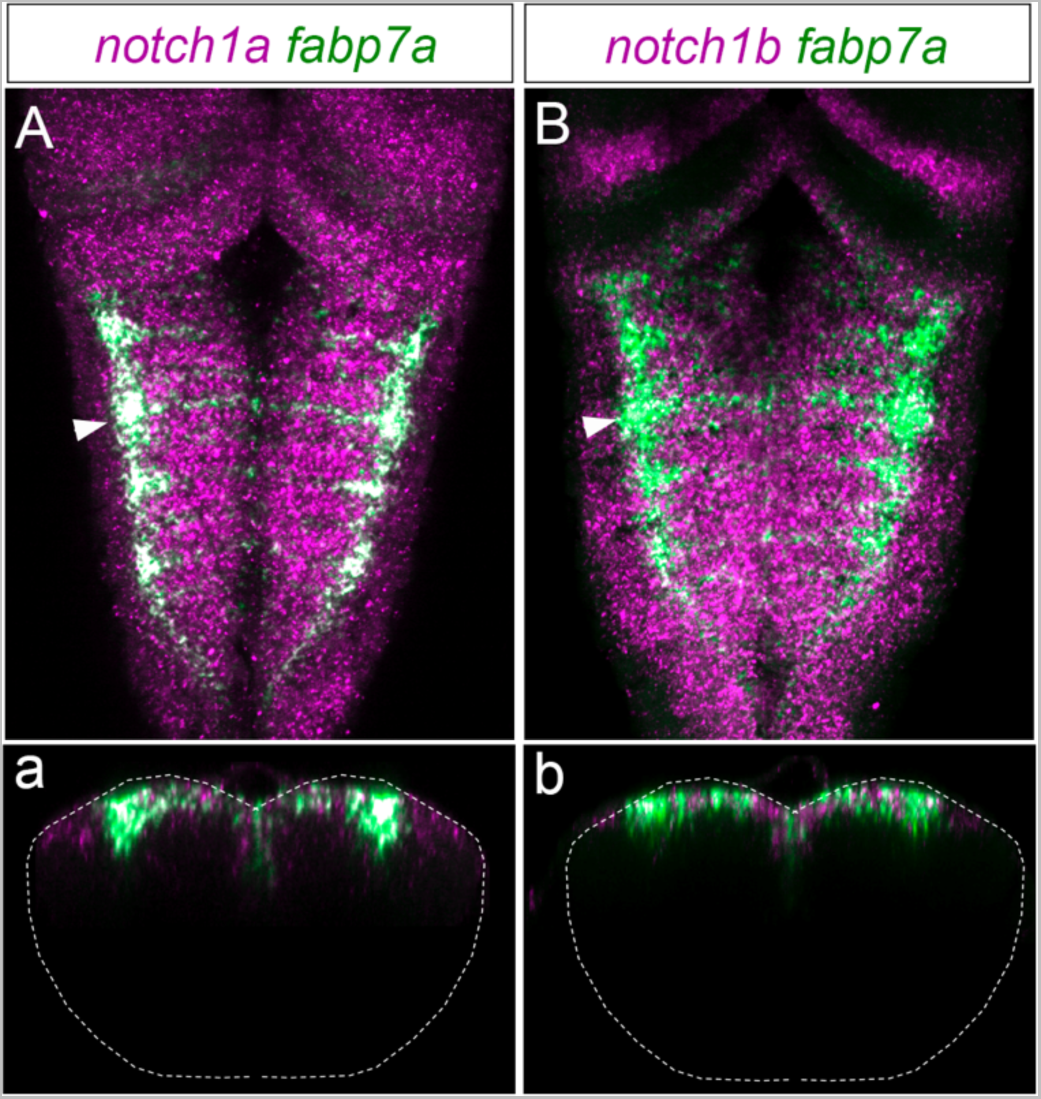
*notch1a* and *notch1b* receptors are expressed all along the AP axis. (A—B) Embryos were *in situ* hybridized with *fabp7a* (green) and *notch1a* or *notch1b* (magenta) at 48hpf. Dorsal MIP with anterior to the top. (a—b) Transverse views of (A—B) through the center of the rhombomere 4 (see white arrowhead).

**Supplementary Figure 2:**
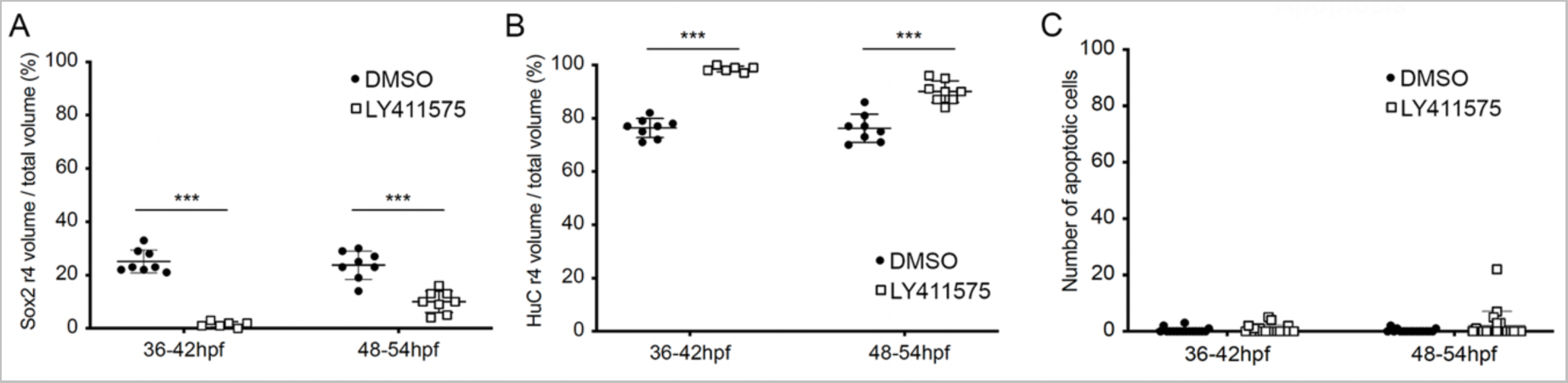
The neuronal differentiation domain increases at the expense of the progenitor domain in the rhombomeres upon inhibition of the Notch-pathway. (A—C) Dot-plots showing the volume of the Sox2 progenitor domain (A), the HuC differentiated domain (B) in r4, and the total hindbrain apoptotic events (C) in embryos treated with DMSO or LY411575 at the indicated intervals. Each dot corresponds to an embryo. (A) The Sox2 volumes of r4 in embryos treated between 36 and 42hpf were: DMSO 25.1 ± 4.3 n=8 vs. LY411575 1.5 ± 1 n=6, p<0.0005; and in embryos treated between 48 and 54hpf were: DMSO 23.7 ± 5.3, n=8 vs. LY411575 10 ± 4.1 n=8, p<0.001. (B) The HuC volumes of r4 in embryos treated between 36 and 42hpf were: DMSO 76.4 ± 3.6 n=8 vs. LY411575 98.5 ± 1 n=6, p<0.0005; and in embryos treated between 48 and 54hpf were: DMSO 75.3 ± 5.3 n=8 vs. LY411575 90 ± 4.1 n=8, p<0.0005. Note the increase of the neuronal differentiation domain accompanied with the decrease of the progenitor domain upon Notch-inhibition. (C) The apoptotic events in embryos treated between 36 and 42hpf were: DMSO 0.3 ± 0.9 n=17 vs. LY411575 0.8 ± 1.5 n=19, ns; and in embryos treated between 48 and 54hpf were: DMSO 0.25 ± 0.6 n=16 vs. LY411575 1.1 ± 2.1 n=17, ns. In all cases the non-parametric Mann-Whitney test was used.

## Notes

### Competing Interest Statement

The authors have declared no competing interest.

## BIBLIOGRAPHY

Allende, M. L., and Weinberg, E. S. (1994). The expression pattern of two zebrafish achaete-scute homolog (ash) genes is altered in the embryonic brain of the cyclops mutant. Developmental Biology 166, 509–530. doi: 10.1006/dbio.1994.1334.

Alunni, A., Krecsmarik, M., Bosco, A., Galant, S., Pan, L., Moens, C. B., et al. (2013). Notch3 signaling gates cell cycle entry and limits neural stem cell amplification in the adult pallium. Development 140, 3335–3347. doi: 10.1242/dev.095018.

Belmonte-Mateos, C., and Pujades, C. (2022). From Cell States to Cell Fates: How Cell Proliferation and Neuronal Differentiation Are Coordinated During Embryonic Development. Front Neurosci-switz 15, 781160. doi: 10.3389/fnins.2021.781160.

Belzunce, I., Belmonte-Mateos, C., and Pujades, C. (2020). The interplay of atoh1 genes in the lower rhombic lip during hindbrain morphogenesis. PLoS ONE 15, e0228225. doi: 10.1371/journal.pone.0228225.

Bertrand, N., Castro, D. S., and Guillemot, F. (2002). Proneural genes and the specification of neural cell types. Nature Reviews Neuroscience 3, 517–530. doi: 10.1038/nrn874.

Bouldin, C. M., and Kimelman, D. (2014). Dual Fucci: A New Transgenic Line for Studying the Cell Cycle from Embryos to Adults. Zebrafish 11, 182–183. doi: 10.1089/zeb.2014.0986.

Brockway, N. L., Cook, Z. T., O’Gallagher, M. J., Tobias, Z. J. C., Gedi, M., Carey, K. M., et al. (2019). Multicolor lineage tracing using in vivo time-lapse imaging reveals coordinated death of clonally related cells in the developing vertebrate brain. Developmental Biology 453, 130–140. doi: 10.1016/j.ydbio.2019.05.006.

Calzolari, S., Terriente, J., and Pujades, C. (2014). Cell segregation in the vertebrate hindbrain relies on actomyosin cables located at the interhombomeric boundaries. The EMBO Journal 33, 686–701. doi: 10.1002/embj.201386003.

Cheng, Y.-C., Amoyel, M., Qiu, X., Jiang, Y.-J., Xu, Q., and Wilkinson, D. G. (2004). Notch activation regulates the segregation and differentiation of rhombomere boundary cells in the zebrafish hindbrain. Developmental Cell 6, 539–550.

Clark, B. S., Cui, S., Miesfeld, J. B., Klezovitch, O., Vasioukhin, V., and Link, B. A. (2012). Loss of Llgl1 in retinal neuroepithelia reveals links between apical domain size, Notch activity and neurogenesis. Development 139, 1599–1610. doi: 10.1242/dev.078097.

Coolen, M., Thieffry, D., Drivenes, Ø., Becker, T. S., and Bally-Cuif, L. (2012). miR-9 controls the timing of neurogenesis through the direct inhibition of antagonistic factors. Developmental Cell 22, 1052–1064. doi: 10.1016/j.devcel.2012.03.003.

Distel, M., Wullimann, M. F., and Köster, R. W. (2009). Optimized Gal4 genetics for permanent gene expression mapping in zebrafish. Proceedings of the National Academy of Sciences of the United States of America 106, 13365–13370. doi: 10.1073/pnas.0903060106.

Dyer, C., Linker, C., Graham, A., and Knight, R. (2014). Specification of sensory neurons occurs through diverse developmental programs functioning in the brain and spinal cord. Dev. Dyn. 243, 1429–1439. doi: 10.1002/dvdy.24184.

Engel-Pizcueta, C., and Pujades, C. (2021). Interplay Between Notch and YAP/TAZ Pathways in the Regulation of Cell Fate During Embryo Development. Frontiers Cell Dev Biology 9, 711531. doi: 10.3389/fcell.2021.711531.

Esain, V., Postlethwait, J. H., Charnay, P., and Ghislain, J. (2010). FGF-receptor signalling controls neural cell diversity in the zebrafish hindbrain by regulating olig2 and sox9. Development 137, 33–42. doi: 10.1242/dev.038026.

Fraser, S., Keynes, R., and Lumsden, A. (1990). Segmentation in the chick embryo hindbrain is defined by cell lineage restrictions. Nature 344, 431–435. doi: 10.1038/344431a0.

Furutachi, S., Miya, H., Watanabe, T., Kawai, H., Yamasaki, N., Harada, Y., et al. (2015). Slowly dividing neural progenitors are an embryonic origin of adult neural stem cells. Nat. Neurosci. 18, 657–665. doi: 10.1038/nn.3989.

Gonzalez-Quevedo, R., Lee, Y., Poss, K. D., and Wilkinson, D. G. (2010). Neuronal regulation of the spatial patterning of neurogenesis. Developmental Cell 18, 136–147. doi: 10.1016/j.devcel.2009.11.010.

Gutzman, J. H., and Sive, H. (2010). Epithelial relaxation mediated by the myosin phosphatase regulator Mypt1 is required for brain ventricle lumen expansion and hindbrain morphogenesis. Development 137, 795–804. doi: 10.1242/dev.042705.

Harada, Y., Yamada, M., Imayoshi, I., Kageyama, R., Suzuki, Y., Kuniya, T., et al. (2021). Cell cycle arrest determines adult neural stem cell ontogeny by an embryonic Notch-nonoscillatory Hey1 module. Nat. Commun. 12, 6562. doi: 10.1038/s41467-021-26605-0.

Hartfuss, E., Galli, R., Heins, N., and Götz, M. (2001). Characterization of CNS Precursor Subtypes and Radial Glia. Dev Biol 229, 15–30. doi: 10.1006/dbio.2000.9962.

Hevia, C. F., Engel-Pizcueta, C., Udina, F., and Pujades, C. (2022). The neurogenic fate of the hindbrain boundaries relies on Notch3-dependent asymmetric cell divisions. Cell Reports 39, 110915. doi: 10.1016/j.celrep.2022.110915.

Itoh, M., and Chitnis, A. B. (2001). Expression of proneural and neurogenic genes in the zebrafish lateral line primordium correlates with selection of hair cell fate in neuromasts. MECHANISMS OF DEVELOPMENT 102, 263–266. doi: 10.1016/s0925-4773(01)00308-2.

Jimenez-Guri, E., Udina, F., Colas, J.-F., Sharpe, J., Padrón-Barthe, L., Torres, M., et al. (2010). Clonal analysis in mice underlines the importance of rhombomeric boundaries in cell movement restriction during hindbrain segmentation. PLoS ONE 5, e10112. doi: 10.1371/journal.pone.0010112.

Krumlauf, R., and Wilkinson, D. G. (2021). Segmentation and patterning of the vertebrate hindbrain. Development. doi: 10.1242/dev.186460.

Labalette, C., Bouchoucha, Y. X., Wassef, M. A., Gongal, P. A., Men, J. L., Becker, T., et al. (2011). Hindbrain patterning requires fine-tuning of early krox20 transcription by Sprouty 4. Development 138, 317–326. doi: 10.1242/dev.057299.

Lam, C. S., März, M., and Strähle, U. (2009). gfap and nestin reporter lines reveal characteristics of neural progenitors in the adult zebrafish brain. Dev Dynam 238, 475–486. doi: 10.1002/dvdy.21853.

Letelier, J., Terriente, J., Belzunce, I., Voltes, A., Undurraga, C. A., Polvillo, R., et al. (2018). Evolutionary emergence of the rac3b/rfng/sgca regulatory cluster refined mechanisms for hindbrain boundaries formation. Proceedings of the National Academy of Sciences of the United States of America 115. doi: 10.1073/pnas.1719885115.

Lowery, L. A., and Sive, H. (2004). Strategies of vertebrate neurulation and a re-evaluation of teleost neural tube formation. MECHANISMS OF DEVELOPMENT 121, 1189–1197. doi: 10.1016/j.mod.2004.04.022.

Lumsden, A. (2004). Segmentation and compartition in the early avian hindbrain. MECHANISMS OF DEVELOPMENT 121, 1081–1088. doi: 10.1016/j.mod.2004.04.018.

Lyons, D. A., Guy, A. T., and Clarke, J. D. W. (2003). Monitoring neural progenitor fate through multiple rounds of division in an intact vertebrate brain. Development 130, 3427–3436. doi: 10.1242/dev.00569.

Nikolaou, N., Watanabe-Asaka, T., Gerety, S., Distel, M., Köster, R. W., and Wilkinson, D. G. (2009). Lunatic fringe promotes the lateral inhibition of neurogenesis. Development 136, 2523–2533. doi: 10.1242/dev.034736.

Park, H. C., Kim, C. H., Bae, Y. K., Yeo, S. Y., Kim, S. H., Hong, S. K., et al. (2000). Analysis of upstream elements in the HuC promoter leads to the establishment of transgenic zebrafish with fluorescent neurons. Developmental Biology 227, 279–293. doi: 10.1006/dbio.2000.9898.

Park, S.-H., Yeo, S.-Y., Yoo, K.-W., Hong, S.-K., Lee, S., Rhee, M., et al. (2003). Zath3, a neural basic helix-loop-helix gene, regulates early neurogenesis in the zebrafish. BIOCHEMICAL AND BIOPHYSICAL RESEARCH COMMUNICATIONS 308, 184–190. doi: 10.1016/s0006-291x(03)01353-6.

Peretz, Y., Eren, N., Kohl, A., Hen, G., Yaniv, K., Weisinger, K., et al. (2016). A new role of hindbrain boundaries as pools of neural stem/progenitor cells regulated by Sox2. BMC biology 14, 57. doi: 10.1186/s12915-016-0277-y.

Pujades, C. (2020). The multiple functions of hindbrain boundary cells: Tinkering boundaries? Seminars in Cell and Developmental Biology 107, 179–189. doi: 10.1016/j.semcdb.2020.05.002.

Riley, B. B., Chiang, M.-Y., Storch, E. M., Heck, R., Buckles, G. R., and Lekven, A. C. (2004). Rhombomere boundaries are Wnt signaling centers that regulate metameric patterning in the zebrafish hindbrain. Developmental Dynamics 231, 278–291. doi: 10.1002/dvdy.20133.

Sahu, A., Devi, S., Jui, J., and Goldman, D. (2021). Notch signaling via Hey1 and Id2b regulates Müller glia’s regenerative response to retinal injury. Glia 69, 2882–2898. doi: 10.1002/glia.24075.

Stigloher, C., Chapouton, P., Adolf, B., and Bally-Cuif, L. (2008). Identification of neural progenitor pools by E(Spl) factors in the embryonic and adult brain. Brain Research Bulletin 75, 266–273. doi: 10.1016/j.brainresbull.2007.10.032.

Stolt, C. C., Lommes, P., Sock, E., Chaboissier, M.-C., Schedl, A., and Wegner, M. (2003). The Sox9 transcription factor determines glial fate choice in the developing spinal cord. Genes & Development 17, 1677–1689. doi: 10.1101/gad.259003.

Tambalo, M., Mitter, R., and Wilkinson, D. G. (2020). A single cell transcriptome atlas of the developing zebrafish hindbrain. Development 147, dev184143. doi: 10.1242/dev.184143.

Than-Trong, E., and Bally-Cuif, L. (2015). Radial glia and neural progenitors in the adult zebrafish central nervous system. Glia 63, 1406–1428. doi: 10.1002/glia.22856.

Than-Trong, E., Ortica-Gatti, S., Mella, S., Nepal, C., Alunni, A., and Bally-Cuif, L. (2018). Neural stem cell quiescence and stemness are molecularly distinct outputs of the Notch3 signalling cascade in the vertebrate adult brain. Development 145. doi: 10.1242/dev.161034.

Thisse, C., and Thisse, B. (2008). High-resolution in situ hybridization to whole-mount zebrafish embryos. Nature Protocols 3, 59–69. doi: 10.1038/nprot.2007.514.

Urbán, N., Blomfield, I. M., and Guillemot, F. (2019). Quiescence of Adult Mammalian Neural Stem Cells: A Highly Regulated Rest. Neuron 104, 834–848. doi: 10.1016/j.neuron.2019.09.026.

Voltes, A., Hevia, C. F., Engel-Pizcueta, C., Dingare, C., Calzolari, S., Terriente, J., et al. (2019). Yap/Taz-TEAD activity links mechanical cues to progenitor cell behavior during zebrafish hindbrain segmentation. Development 146. doi: 10.1242/dev.176735.

